# Triclabendazole inhibits succinate dehydrogenase *in vitro*

**DOI:** 10.1101/2025.11.20.689426

**Authors:** Tobias Kämpfer, María Eugenia Ancarola, Pascal Zumstein, Alice Bernal, Trix Zumkehr, Seraina Mühlemann, Natalie Wiedemar, Britta Lundström-Stadelmann

**Affiliations:** Institute of Parasitology, Department of Infectious Diseases and Pathobiology, Vetsuisse Faculty, University of Bern, Länggassstrasse 122, 3012 Bern, Switzerland; Graduate School for Cellular and Biomedical Sciences (GCB), University of Bern, Mittelstrasse 43, 3012 Bern, Switzerland; Multidisciplinary Center for Infectious Diseases, University of Bern, Hallerstrasse 6, 3012 Bern, Switzerland

**Keywords:** succinate dehydrogenase, triclabendazole, parasite, helminth, anthelmintic, complex II, benzimidazole, *Echinococcus*

## Abstract

Succinate dehydrogenase (SDH, complex II), is essential for mitochondrial respiration. This study shows that triclabendazole and its primary metabolites inhibit SDH activity in mitochondria of the cestode *Echinococcus multilocularis*, while other tested benzimidazoles do not. Further analyses revealed that triclabendazole and metabolites also inhibit SDH in mitochondria of additional helminths (trematodes, nematodes) and mammalian cells. These findings indicate a shared mode of action and highlight potential safety risks associated with long-term triclabendazole treatment in mammals.

## Manuscript text

In eukaryotic organisms, energy generation primarily relies on mitochondrial oxidative phosphorylation, which integrates the electron transport chain (ETC) and substrate-level phosphorylation. Electrons pass through enzyme complexes (I-IV) to the final acceptor, oxygen. This creates a proton gradient across the inner mitochondrial membrane that powers ATP-synthase (complex V) (1). As many parasites depend on mitochondrial energy metabolism during specific life cycle stages and their mitochondrial components often differ from those of mammalian hosts, the ETC has emerged as a selective drug: Several antiparasitic agents (2–5) act on complex III (cytochrome *bc*_1_), leading to rapid energy depletion and parasite death. Another promising target is mitochondrial complex II, which acts as a succinate dehydrogenase (SDH). By oxidizing succinate to fumarate, it links the citric acid cycle with oxidative phosphorylation. In helminths and other organisms living under low-oxygen conditions, this enzyme can catalyze the reverse reaction, acting as a fumarate reductase with fumarate as the terminal electron acceptor (6). It is essential for the energy metabolism of many helminths and represents an attractive, potentially druggable, target (7, 8).

*E. multilocularis* causes alveolar echinococcosis, a severe zoonosis with infiltrative, tumor-like growth of larval cysts (metacestodes) in the liver and fatal outcome if untreated (9). Drug treatment options are limited to the benzimidazoles (BMZs) albendazole and mebendazole, which are parasitostatic, often cause side effects, and rarely achieve cure (10), highlighting the need for safer and more effective drugs. BMZs inhibit microtubule polymerization (11), but may also target other pathways in helminths (12), including mitochondrial complex II (13–15). To explore this, we screened 20 BMZs (Table 1) for activity on SDH in mitochondria from *E. multilocularis* metacestode vesicles, other helminths (trematodes and nematodes), and mammalian cells.

**Table 1.** List of tested benzimidazoles.

| Benzimidazole | Abbreviation | Supplier |
| --- | --- | --- |
| Albendazole | ABZ | Merck, Buchs, Switzerland |
| Ricobendazole | ABZ-SO | Lucerna-Chem, Lucerne, Switzerland |
| Albendazole sulfone | ABZ-SO<sub>2</sub> | Lucerna-Chem, Lucerne, Switzerland |
| Benomyl | BNM | Lucerna-Chem, Lucerne, Switzerland |
| Cambendazole | CMB | Lucerna-Chem, Lucerne, Switzerland |
| Carbendazim | CZM | Lucerna-Chem, Lucerne, Switzerland |
| Fenbendazole | FBZ | Merck, Buchs, Switzerland |
| Febantel | FBT | Merck, Buchs, Switzerland |
| Mebendazole | MBZ | Lucerna-Chem, Lucerne, Switzerland |
| Nocodazole | NCZ | Lucerna-Chem, Lucerne, Switzerland |
| Oxibendazole | OBZ | Lucerna-Chem, Lucerne, Switzerland |
| Oxfendazole | OFZ | Merck, Buchs, Switzerland |
| Omeprazole | OMP | Lucerna-Chem, Lucerne, Switzerland |
| Parbendazole | PBZ | Lucerna-Chem, Lucerne, Switzerland |
| Triclabendazole | TCBZ | Lucerna-Chem, Lucerne, Switzerland |
| Triclabendazole sulfoxide | TCBZ-SO | Merck, Buchs, Switzerland |
| Triclabendazole sulfone | TCBZ-SO<sub>2</sub> | Merck, Buchs, Switzerland |
| Thiabendazole | THB | Lucerna-Chem, Lucerne, Switzerland |
| Flubendazole | UBZ | Lucerna-Chem, Lucerne, Switzerland |

*E. multilocularis* metacestode vesicles were cultured according to Spiliotis et al. (16). Adult *Fasciola hepatica* flukes isolated from the bile ducts of naturally infected cattle livers (obtained from local abattoir in Switzerland), and adult *Toxocara cati* from the diagnostic unit of the Institute of Parasitology, University of Bern, Switzerland, served as trematode and nematode models, respectively. Mammalian controls included mitochondria from rat Reuber hepatoma H-4-II-E cells (RH cells, *Rattus norvegicus*, ATCC, LGC Standards S.a.r.l., Molsheim Cedex, France) and liver tissue from healthy female mice (*Mus musculus*, CD-1 IGS strain, Charles River Laboratories, Germany).

SDH activity was measured fluorometrically via irreversible reduction of resazurin to resorufin (17), preventing reverse complex II activity. As control, an assay with triclabendazole (TCBZ), no mitochondria, and alternative reducing agents was performed (Supplementary Figure 1). Mitochondria were prepared by tissue homogenization in buffer (Tris-HCl 15 mM, pH 7.4, sucrose 0.33 M, EDTA 25 µM) with a glass-Teflon homogenizer, followed by centrifugation at 1’000 x g for 10 min, 4°C. Enriched mitochondrial fractions were diluted to 10 mg/mL in hypotonic buffer (Tris-HCl 10 mM, pH 7.5) and used at 0.1 mg/mL in 100 µL assay buffer (50 mM potassium phosphate pH 7.5, 1 mM MgCl_2_, 5 mM NaN_3_, 27.5 mM resazurin) with either 2% DMSO (negative control), 2% DMSO + 100 mM sodium malonate (positive control), or 2% DMSO + 40 µM BMZ. Reactions were started with 40 mM succinate and measured in 96-well plates with three biological replicates and technical triplicates. Fluorescence (λ_ex_ = 544 nm, λ_em_ = 590 nm) was measured every 30 sec for 5 min before and for 10 min after initiation (Hidex sense reader, Hidex Oy, Turku, Finland). Relative SDH activity was expressed as fluorescence change after succinate addition relative to the negative control.

The BMZs were screened on mitochondria enriched from *E. multilocularis* metacestode vesicles (Figure 1). The positive control (malonate), as well as TCBZ and its metabolites (TCBZ-SO and TCBZ-SO_2_) significantly inhibited SDH activity compared to the DMSO negative control (malonate: 15.0%; TCBZ: 25.9%; TCBZ-SO: 59.2%; TCBZ-SO_2_: 46.3%; see Supplementary File 2 for data and respective *p*-values). In contrast, all other BMZs showed SDH activity above 80%.

**Fig. 1.**
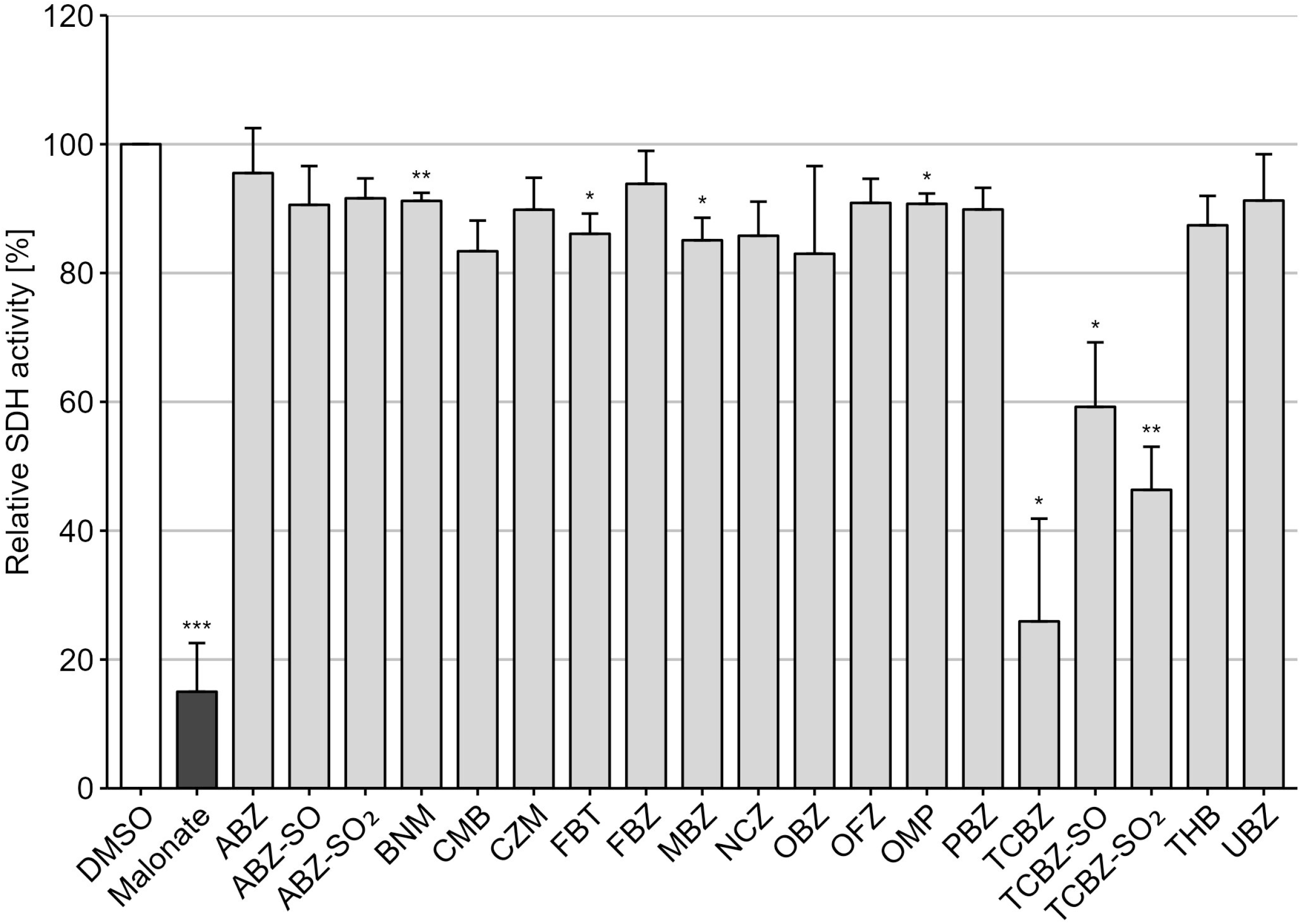
SDH activity assays performed on enriched mitochondrial fractions from *E. multilocularis* metacestode vesicles. Measurements were performed in biological and technical triplicates. Data is shown as mean fluorescence increase relative to the negative control +SD. Results with Bonferroni-adjusted *p*-values below 0.05 (*), 0.01 (**), and 0.001 (***) were regarded as significant.

To further explore this observation, we tested TCBZ, albendazole (ABZ), and their respective metabolites on mitochondria-enriched samples from other organisms (Figure 2). TCBZ, its primary metabolites, and the malonate control, consistently inhibited SDH activity across all tested helminth species and mammalian models. Significant inhibition was observed in *F. hepatica* (TCBZ: 3.08% TCBZ-SO: 55.0%; TCBZ-SO_2_: 46.6%), *T. cati* (TCBZ: 52.3%; TCBZ-SO: 51.3%; TCBZ-SO_2_: 48.4%), *R. norvegicus* (TCBZ: 45.8%; TCBZ-SO: 36.6%; TCBZ-SO_2_: 26.8%), and *M. musculus* (TCBZ: 46.4%; TCBZ-SO: 66.1%; TCBZ-SO_2_: 39.7%; see Supplementary File 2 for data and respective *p*-values). In contrast, ABZ and its metabolites showed no significant effect on SDH activity.

**Fig. 2.**
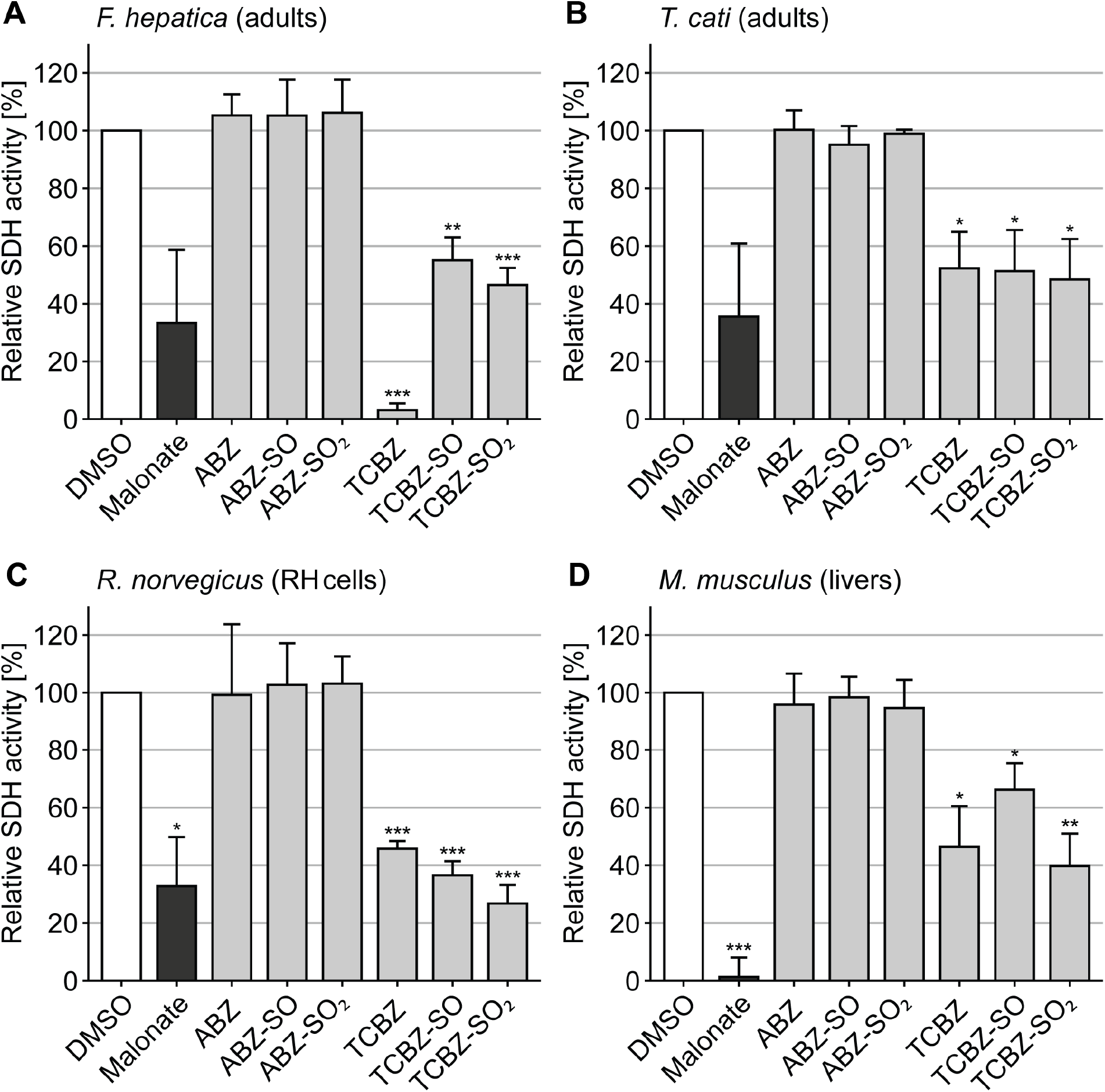
SDH activity assay performed on enriched mitochondrial fractions from adult flukes of *F. hepatica* (A), adult worms of *T. cati* (B), cultured hepatoma cells of *R. norvegicus* (C), and livers of mice (*M. musculus*) (D). Measurements were performed in biological and technical triplicates. Data is shown as mean fluorescence increase relative to the negative control +SD. Results with Bonferroni-adjusted *p*-values below 0.05 (*), 0.01 (**), and 0.001 (***) were regarded as significant.

**Fig. 3.**
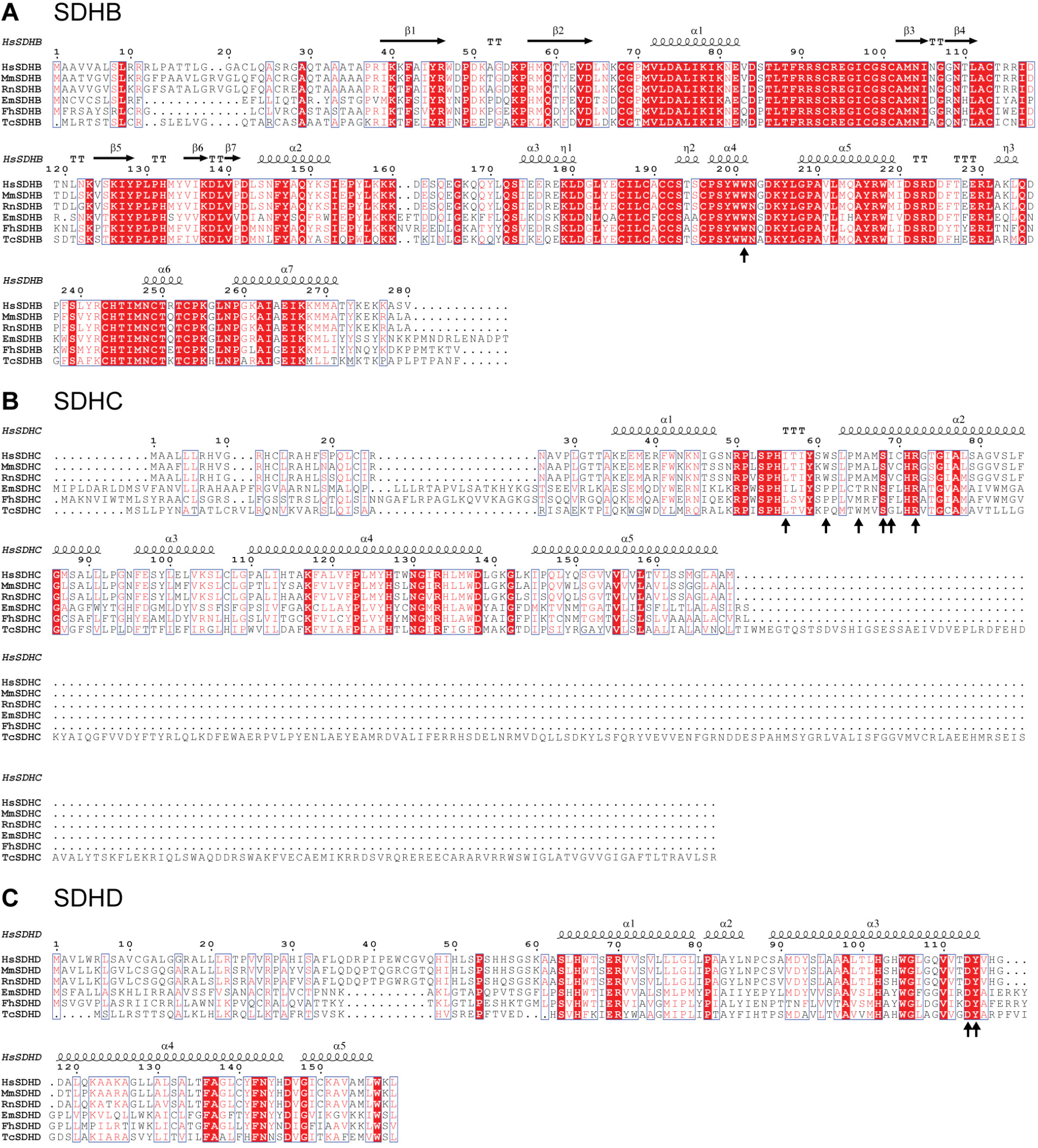
Multiple sequence alignment of SDH subunits (SDHB: succinate dehydrogenase [quinone] iron-sulfur subunit, SDHC: succinate dehydrogenase cytochrome b560 subunit, SDHD: succinate dehydrogenase [quinone] cytochrome b small subunit). Red indicates identical residues, while red letters indicate similar residues. Black arrows indicate the quinone-binding site according to Inaoka et al., 2015 (28). As *T. cati* sequences were unavailable, *T. canis* sequences were used instead.

These findings show that TCBZ targets complex II in a broad, species-independent manner across all tested helminth groups and also in mammals. This contrasts with earlier studies, which primarily focused on nematodes (18–23).

To assess potential structural explanations, we compared SDH subunit sequences using BioEdit (v7.2), with visualization in ESPript 3.0. Sequences were retrieved from UniProt (Supplementary File 3), with secondary structure information from the human complex II (PDB:8GS8). Attention was given to subunits B-D, which form the quinone-binding site, a described target of thiabendazole (13, 24). Alignments for subunit A are shown in Supplementary Figure 4. Most subunits were conserved, but distinct residue changes appeared in the quinone-binding site of helminths compared with mammals, consistent with their use of alternative quinones such as rhodoquinone (25, 26).

TCBZ is currently used to treat *F. hepatica* infections, usually as a well-tolerated single dose (27). In contrast, treatment of tissue-dwelling parasites like *E. multilocularis* require prolonged, multi-dosage regimens with other BMZs (10). Based on our findings, we advise caution regarding the long-term application of TCBZ in mammalian hosts, as toxic effects on mitochondrial function may occur. Nevertheless, our data highlights TCBZ as a promising drug for the development of new therapies against life-threatening helminth infections such as alveolar echinococcosis and other zoonotic helminthiases.

## Acknowledgements

We would like to thank the diagnostic unit of the Institute of Parasitology, Bern for kindly providing *Toxocara* material. We hank Kai Hänggeli from the Institute of Parasitology, University of Bern for supporting us in mouse material processing. We would also like to thank Matías Preza from the Institute Parasitology, Bern, for his input in the establishment of the enzymatic assays applied in this study.

## Supplemental Material

Supplementary figure 1: Resazurin reduction confirmation assay showing that TCBZ does not inhibit resazurin reduction in the absence mitochondrial dehydrogenases.

Supplementary file 2: Data of SDH inhibition assay and respective *p*-values.

Supplementary file 3: List of UniProt accession numbers of the sequences used for the sequence alignment.

Supplementary figure 4: Sequence alignments for subunit A (succinate dehydrogenase [quinone] flavoprotein subunit) of mitochondrial complex II.

## Data availability

The data presented in the manuscript is available in the supplementary material. The authors confirm that all protocols are detailed within this article.

## Author Contributions

Conceptualization TK, BLS

Data curation TK, MEA

Formal analysis TK, MEA, PZ

Funding Acquisition BLS, NW

Investigation TK, MEA, PZ, AB, SM

Methodology TK, MEA, PZ, AB

Project Administration BLS

Resources BLS, NW

Supervision BLS, NW

Validation MEA, TZ, SM

Visualization TK, MEA, PZ

Writing – original draft TK, BLS Writing – review & editing all authors

## Ethics Statement

Animals used in this study for parasite strain maintenance and cell extracts were kept in compliance with the Swiss animal protection law (TschV, SR 455) and experiments were approved by the Animal Welfare Committee of the Canton of Bern (license numbers BE2/22 and BE40/21). Animals were housed in type 3 cages, enriched with a plastic house, paper and woodchip bedding. They were maintained in a 12 h light/dark cycle, controlled temperature of 21 °C – 23 °C, and a relative humidity of 45% - 55%. Food and water were provided *ad libitum*.

